# C*is* non-coding genetic variation drives gene expression changes in the *E. coli* and *P. aeruginosa* pangenomes

**DOI:** 10.1101/2025.06.10.658842

**Authors:** Bamu F. Damaris, Matylda Zietek, Jelena Erdmann, Athanasios Typas, Susanne Häußler, Marco Galardini

**Affiliations:** Institute for Molecular Bacteriology, TWINCORE Centre for Experimental and Clinical Infection Research, a joint venture between the Hannover Medical School (MHH) and the Helmholtz Centre for Infection Research (HZI), Hannover, Germany; Cluster of Excellence RESIST (EXC 2155), Hannover Medical School (MHH), Hannover, Germany; Genome Biology Unit, European Molecular Biology Laboratory (EMBL), Heidelberg, Germany; Molecular Systems Biology Unit, EMBL, Heidelberg, Germany; Department of Clinical Microbiology, Copenhagen University Hospital - Rigshospitalet, Copenhagen, Denmark

## Abstract

Bacteria use gene regulation to dynamically adapt to changes in their environment, including resistance to stress and the occupation of new niches. Gene expression is known to vary within a species pangenome, but the extent to which these changes could be explained by genetic variants in *cis* non-coding regions has so far been poorly investigated. Statistical genetics offers a hypothesis-free approach to this problem, as opposed to mechanistic models, which can be used only for reference isolates that are not representative of the whole species. In this study, we assembled two genomic and transcriptomic datasets for *Escherichia coli* (N=117) and *Pseudomonas aeruginosa* (N=413) and identified associations between genetic variants in *cis* non-coding regions and recorded gene expression variation. We identified at least one associated variant in up to 39% of the tested genes in both species. We partly validated the associations *in-silico* and *in-vitro* for *E. coli*, reinforcing the difficulty of identifying a single mechanism generating gene expression diversity. We then investigated the relevance of non-coding variants in explaining the variability in antimicrobial resistance in both species using two additional publicly available datasets, identifying a large number of these variants across antimicrobial compounds. This work confirms the role of genetic variation in often overlooked regions of bacterial genomes in influencing molecular and clinically relevant phenotypes.

## Introduction

Population genetics principles^1^, harsh competition, and the generally limited resource settings faced by prokaryotes suggest that microbial cells will manage their resources efficiently, therefore tending to express some of their genes only when necessary for survival^2^. Examples of this general principle have been known in the *E. coli* carbon metabolism since the early days of molecular microbiology^3^. This principle also applies to survival to stresses, such as antimicrobial resistance (AMR), as well as occupying new ecological niches, such as the human host for pathogenic species^4^.

The same efficiency principle results in physically interacting and functionally related genes being encoded in operons, usually controlled by a single promoter. The expression of whole functional units encoded in a prokaryotic cell could then largely depend on regulatory elements located in the non-coding region immediately upstream of an operon (*cis* regulation). These non-coding regions contain regulatory elements such as canonical promoters recognized by specific sigma-factors (σ) of the RNA-polymerase, as well as transcription factor binding sites (TFBS) that determine the binding of transcription factors, which in turn favour (activators) or repress (repressors) gene expression^5^. These non-coding regions have in fact been shown to be under selection, suggesting that they are evolutionarily constrained because of the impact that mutations in these regions could ultimately have on survival^6,7^. Given the high plasticity of bacterial genomes, genetic variants in non-coding regions can take up multiple forms, each of which may alter gene expression. Short variants such as SNPs/InDels have been shown to be able to create active promoter sequences from random sequences when put under selection^8^. Larger genetic variants such as promoter switching enable genes to be wired to a different section of the regulatory network, thus enabling phenotypic changes that do not depend on altering the coding sequence^9^.

Even though the field of synthetic biology has over the years accumulated knowledge on how to combine the presence of specific regulatory elements to achieve the desired level of expression, how much natural genetic variation influences observed variations across different bacterial isolates is currently unknown. Although non-coding regions only account for 10-15% of a prokaryotic genome on average, the impact of genetic variants in those regions could result in large phenotypic changes, comparable to those observed for variants in coding regions, as well as gene presence/absence patterns. These changes could then affect observable phenotypes such as antimicrobial resistance^10^ and pathogenicity^11^.

Correctly predicting the impact of non-coding variants on gene expression is a difficult task due to the large number of regulatory elements contributing to the outcome and how they interact with each other, not all of which have been studied. Detailed biophysical models and large scale library screens have been able to improve the understanding of the relationship between primary sequences and corresponding gene expression levels^12,13^, but these are limited to the reference laboratory isolate of *E. coli*, and cannot be used to generalize to natural isolates and across genes and species.

In this study we set out to identify non-coding genetic variants affecting gene expression in a collection of *E. coli* and *Pseudomonas aeruginosa* isolates, for which we obtained both genomic and transcriptomic data. We used a statistical genetics approach that does not rely on prior knowledge of regulatory elements, and only used the assumption that genetic variants affecting gene expression would be located in non-coding regions located in *cis*. In order to capture all sources of genetic variation affecting gene expression levels, we encoded genetic variants using traditional short variants (SNPs/InDels), promoter switching, and gene cluster k-mers^14^.

We could find at least one *cis* genetic variant associated with gene expression across isolates for up to 39% of the genes we tested (713 genes for *E. coli* and 1,632 genes in *P. aeruginosa*), some of which we validated experimentally and could mechanistically explain in *E. coli* thanks to the detailed knowledge of its canonical promoter structure and TFBS sequence motifs. We then showed how antimicrobial resistance is associated with non-coding variants in both species, and validated it using transcriptomics under tobramycin treatment. Our study provides an approach for identifying genetic variants affecting gene expression and resulting phenotypes in ever larger genomic and transcriptomic bacterial collections.

## Results

### Natural and clinical isolates of *E. coli* and *P. aeruginosa* exhibit distinct gene expression profiles compared to reference strains

In this study we aimed to map the role of *cis* non-coding genetic variants in the regulation of gene expression in *E. coli* and *P. aeruginosa*, using an hypothesis-free approach. We analyzed transcriptomic profiles in rich media and genomic data from 117 *E. coli* and 413 *P. aeruginosa* isolates (**Figure 1**). The *E. coli* isolates were mostly clinical (45%) and natural isolates (30%), with lab strains (5%) and experimental evolution isolates (20%) being the minority. The *P. aeruginosa* isolates were all clinical isolates. We measured the transcriptomes of *E. coli* as part of this study, and obtained transcriptomic information for *P. aeruginosa* isolates from a previous study^10,15^. Through a differential gene expression analysis between each isolate and their respective reference genome we revealed significant differences in gene expression among the isolates. To account for differences in gene presence/absence patterns in the species’ pangenome, we defined differentially expressed genes (DEGs) as those present in at least 10% of isolates, with an absolute log_2_ fold-change ≥ 2 in at least 5% of isolates, and an adjusted p-value < 0.0001.

**Figure 1:**
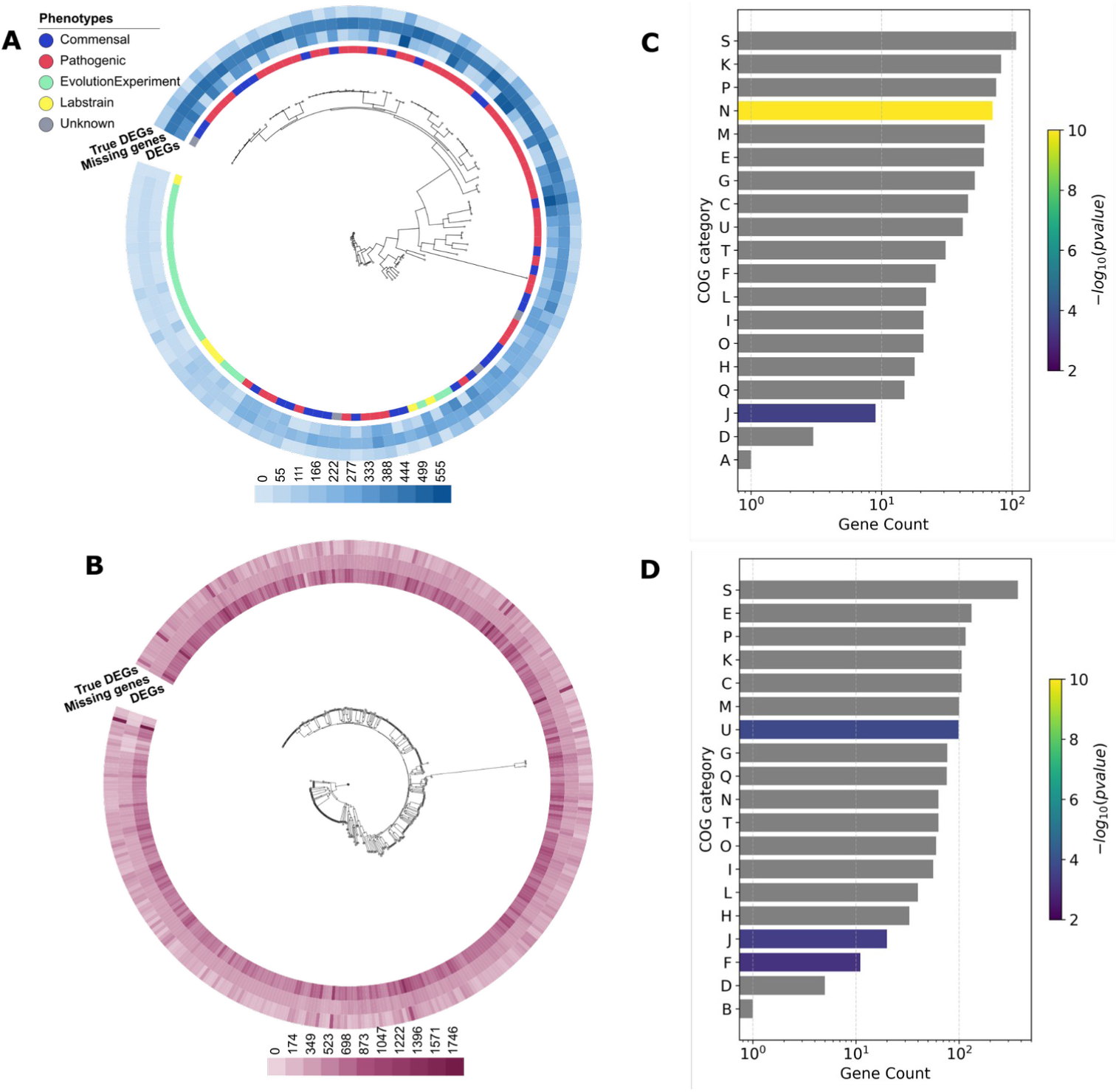
Variability in gene expression in a panel of *E. coli* and *P. aeruginosa* strains. **(A-B)** Phylogenetic trees based on the core genomes of the *E. coli* **(A)** and *P. aeruginosa* **(B)** isolates. The color intensity of the outer rings reflects the gene count within each category. “Missing genes” refers to those absent in the genome of the corresponding isolate in the tree. DEGs are those with an absolute log_2_ fold-change ≥ 2 and an adjusted p-value < 0.0001. “True DEGs” are the number of DEGs excluding those that are missing from the genome of the corresponding isolate in the tree. **Panels (C)** and (**D)** illustrate the functional diversity of DEGs in *E. coli* and *P. aeruginosa* isolates, respectively, compared to their reference strains. The horizontal bars represent the number of genes associated with each COG category (**Supplementary Table 3**). The color scale reflects the statistical significance adjusted for multiple testing.

In *E. coli*, we identified 713 DEGs accounting for 17% of the K-12 proteome. A sizable portion of these genes were those associated with virulence (19%) and antimicrobial resistance (14%). Most of these genes (66%) were a part of operons with diverse biological functions (**Supplementary Table 1**). For example, all genes within the *fimAICDFGH* operon were differentially expressed. This operon encodes type I fimbriae, which facilitates bacterial adhesion and invasion of host tissues^16^. Similarly, within the *acrAB* operon, *acrB* alone was differentially expressed across strains. This operon encodes for part of the AcrAB/TolC multidrug efflux pump, which contributes to antimicrobial resistance, as well as adaptation to the host intestinal environment by exporting bile salts and enabling bacterial survival in host immune cells^17^. We then performed a functional enrichment analysis on the DEGs, which revealed an enrichment of two major COG (cluster of orthologous groups) categories (**Figure 1C**): cell motility and translation, and ribosomal structure and biogenesis. With the key genes in the cell mobility category being *fliC*, which encodes flagellin protein essential for flagellar filaments, *motA* and *motB*, which are necessary for motor rotation^18^. For the translation and ribosomal structure category, notable genes included *hchA*, a stress-response gene^18^.

Since we had transcriptomic data for more *P. aeruginosa* strains, we uncovered more than double of DEGs (1,632), corresponding to 27% of the PA14 reference proteome. Among these, 10% were associated with virulence, and 4% associated with antimicrobial resistance. A substantial proportion of the DEGs (25%) were part of an operon, many of which contributed to bacterial survival and adaptability (**Supplementary Table 2**). For instance, three of the six genes in the alginate biosynthesis operon (*alg44KEGXL*), which regulates alginate biosynthesis^19^, were differentially expressed. Four out of seven genes in the *pelABCDEFG* operon, which encode polysaccharides critical for environmental persistence and biofilm formation^20^, were differentially expressed. Another operon of note was *flgBCDE*, where two out of four genes were differentially expressed. This operon plays a central role in flagellar assembly and motility^21^. Several DEGs, including *mexAB-oprM*, *mexXY*, and *mexCD-oprJ*, were identified in operons associated with multidrug resistance^22^. The DEGs were enriched in several functional categories (**Figure 1D**), including intracellular trafficking, secretion, and vesicular transport. Key genes in this category include *exbB2*, transmembrane proteins, and *pilQ*/*pilC*, which are essential for the assembly of a type IV pilus^23^. Another prominent category was translation, ribosomal structure, and biogenesis, represented by genes such as *rpmJ* (50s ribosomal protein L36), *rpsT* (30S ribosomal protein S20), and *rpIT* (50S ribosomal protein L20)^18^.

These results highlight the significant transcriptional differences between the isolates and their reference genomes in *E. coli* and *P. aeruginosa*. The observed transcriptional variations are likely driven by genetic variants, some of which might be in *cis* (*i.e.* in direct proximity) with respect to each DEG or operon.

### A large fraction of gene expression changes can be attributed to genetic variations in *cis* non-coding regions

Next, we used three types of genetic markers to characterise the genetic variants within the *cis* non-coding regions of the DEGs, which were: short variants like single nucleotide polymorphisms (SNPs) and short insertions and deletions (InDels), the presence/absence of intergenic regions (IGRs), and k-mers. By using these different ways of encoding variants, we aimed to compensate for the limitations of each encoding method, to capture a larger fraction of variants, and lastly to improve the interpretation for those variants significantly associated with gene expression changes. We identified a total of 136,690 short variants within the non-coding region of DEGs in *E. coli*, with SNPs comprising the vast majority (85%). Similarly, in *P. aeruginosa*, we identified 6,125,235 short non-coding variants in the non-coding region of DEGs, where SNPs accounted for 86% of the total variants. We also analyzed the variability in the presence/absence patterns of entire IGRs. We identified 12,491 IGRs within *E. coli* genes, with 71.6% classified as cloud IGRs, meaning they appeared in 15% or fewer of the genomes analyzed. In *P. aeruginosa*, 22,788 IGRs were identified, with 80.9% being cloud IGRs. A smaller proportion of the core IGRs (found in at least 99% of genomes) was comparable in both species, accounting for 10% in *E. coli* and 10.2% in *P. aeruginosa* (**Supplementary** Figure 1).

We then conducted a GWAS analysis to establish statistical associations between each gene’s *cis* non-coding variants and their corresponding expression. We focused on DEGs that had at least a four-fold expression change when compared to that of the reference strains in ≥ 5% of isolates, thereby prioritizing genes with significant change of gene expression. We observed that short variants were associated with the expression of 7% of the genes in *E. coli* and 25% in *P. aeruginosa* (**Figure 2**). When measuring associations using IGRs presence or absence within DEGs, we observed that 28% of the DEGs in *E. coli* were linked to IGR variability, whereas this was true only for 3% of DEGs in *P. aeruginosa*. We also observed that k-mers encoding genetic variants, which captures both short variants and IGR presence/absence patterns, improved the sensitivity of our association analysis; in fact we identified significant associations for 39% of the DEGs in both species. The underlying non-coding genetic variants we identified were associated with both upregulated and downregulated gene expression. We observed that when these *cis* short non-coding variants were present in the first gene of an operon, they affected the expression of 6% of downstream genes in the operon in *E. coli* and 35% in *P. aeruginosa*. For both species we have therefore identified significant associations between *cis* variants and changes in gene expression for up to 39% of DEGs. Since promoters and transcription-factor binding sites (TFBSs) are within regions of these *cis*-variants, we reasoned that these elements could at least partially explain these associations.

**Figure 2:**
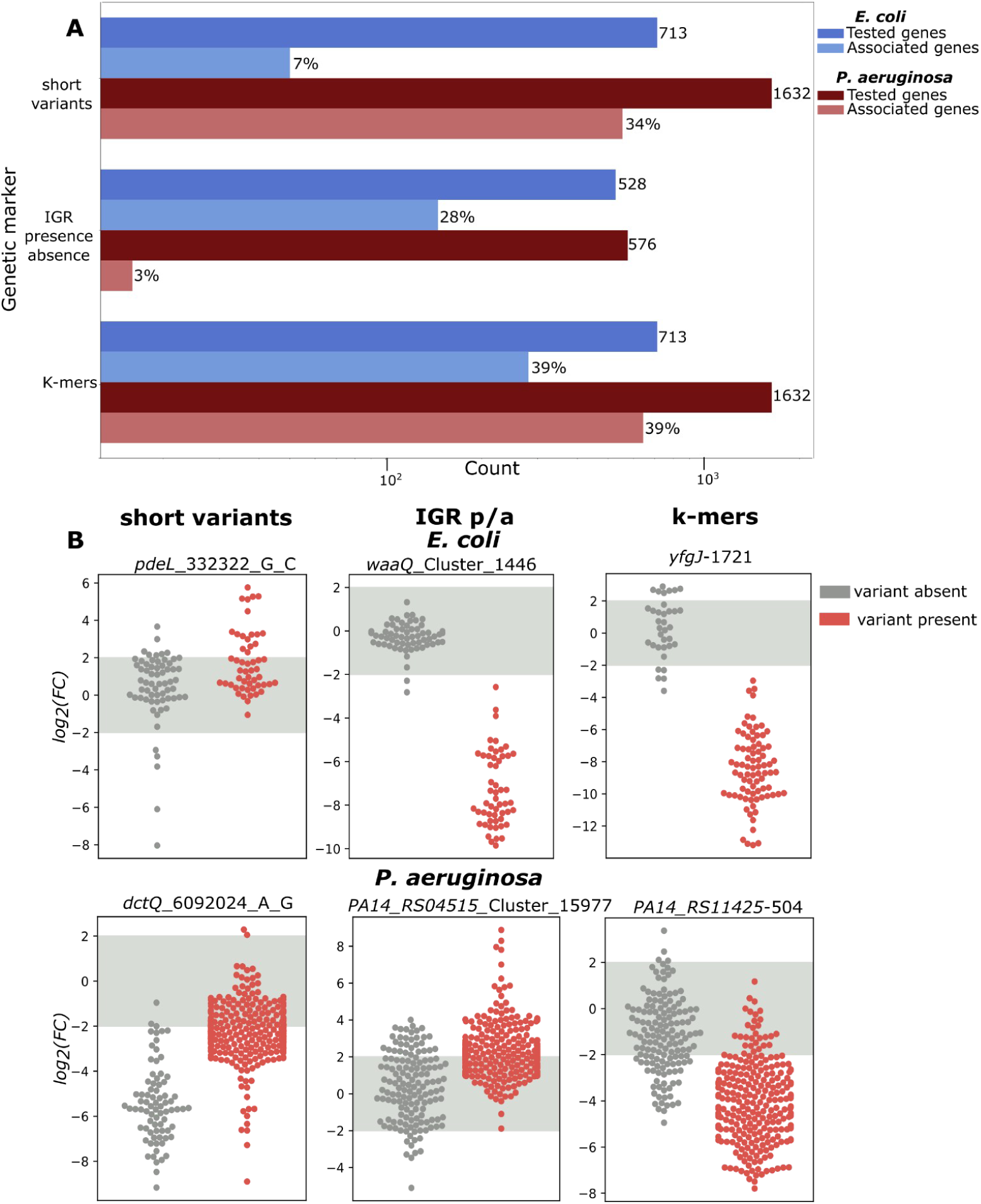
Non-coding variants are associated with gene expression in *E. coli* and *P. aeruginosa*. **(A)** Summary statistics about the relationship between each genetic marker (short variants, IGR presence/absence, and gene cluster specific k-mers) and gene expression in *E. coli* (blue) and *P. aeruginosa* (maroon). **(B)** Each plot shows fold change for each gene associated with SNPs (left), IGR presence/absence (middle), and gene-clustered k-mers (right). Genes include *pdeL*, *waaQ*, and *yfgJ* in *E. coli*, and *dctQ*, *PA14_RS04515*, and *PA14_RS11425* in *P. aeruginosa*. Each point represents an isolate: grey indicates absence of the non-coding variant, red indicates presence. The grey shaded region marks the differential expression threshold; points (*i.e.* isolates) outside this range are considered to have the gene differentially expressed.

### Validation of the effects of non-coding variants on gene expression in *E. coli*

Our association analysis found that variation in gene expression could be explained by the presence of *cis* non-coding variants for up to 39% of DEGs, when encoded using kmers. We then set out to determine the potential mechanism by which these variants exert their effect. We focused these validation approaches for *E. coli*, owing to its well-curated and characterized transcriptional regulatory elements, such as canonical promoters and transcription factors (TFs). To investigate whether non-coding variants previously identified through GWAS are likely to influence gene expression through alterations to canonical promoters, we used a biophysical model^13^, which enabled us to predict transcription rates from the variant-containing non-coding sequences. This model specifically estimates transcription rates based on the presence of putative σ^70^ promoters, the primary sigma factor responsible for housekeeping gene expression under standard conditions. We hypothesized that genetic variants present in the site of a canonical promoter could result in altered transcription rates, and that changes in transcription rates would correlate with the measured gene expression. We focused on the 218 DEGs in *E. coli* whose expression levels were significantly associated with non-coding variants, as identified in our k-mer based GWAS. For each gene, we computed the Pearson’s correlation coefficient between predicted transcription rates (from the promoter calculator) and measured gene expression across the isolates.

We observed that 30% and 3% of DEGs had a moderate (0.2 < |r| < 0.8) or strong (|r| ≥ 0.8) correlation between the predicted transcription rates and the measured gene expression, respectively (**Figure 3A**). We assessed whether these patterns were stronger than what would occur by chance. We performed a permutation, in which the predicted transcription rates were randomly shuffled 1,000 times per gene across isolates and a Pearson’s correlation coefficient was computed between them and the measured gene expression values. We observed that the empirical distribution of correlations coefficients differed significantly from the shuffled ones, using a Kolmogorov-Smirnoff test (p-value < 0.0001). A Q-Q plot comparing the observed and expected correlation values further showed enrichment at the tails of the distribution (**Figure 3B**), further supporting the hypothesis that these DEGs are driven by variants affecting promoters, and that both positive and negative regulatory effects are possible.

**Figure 3:**
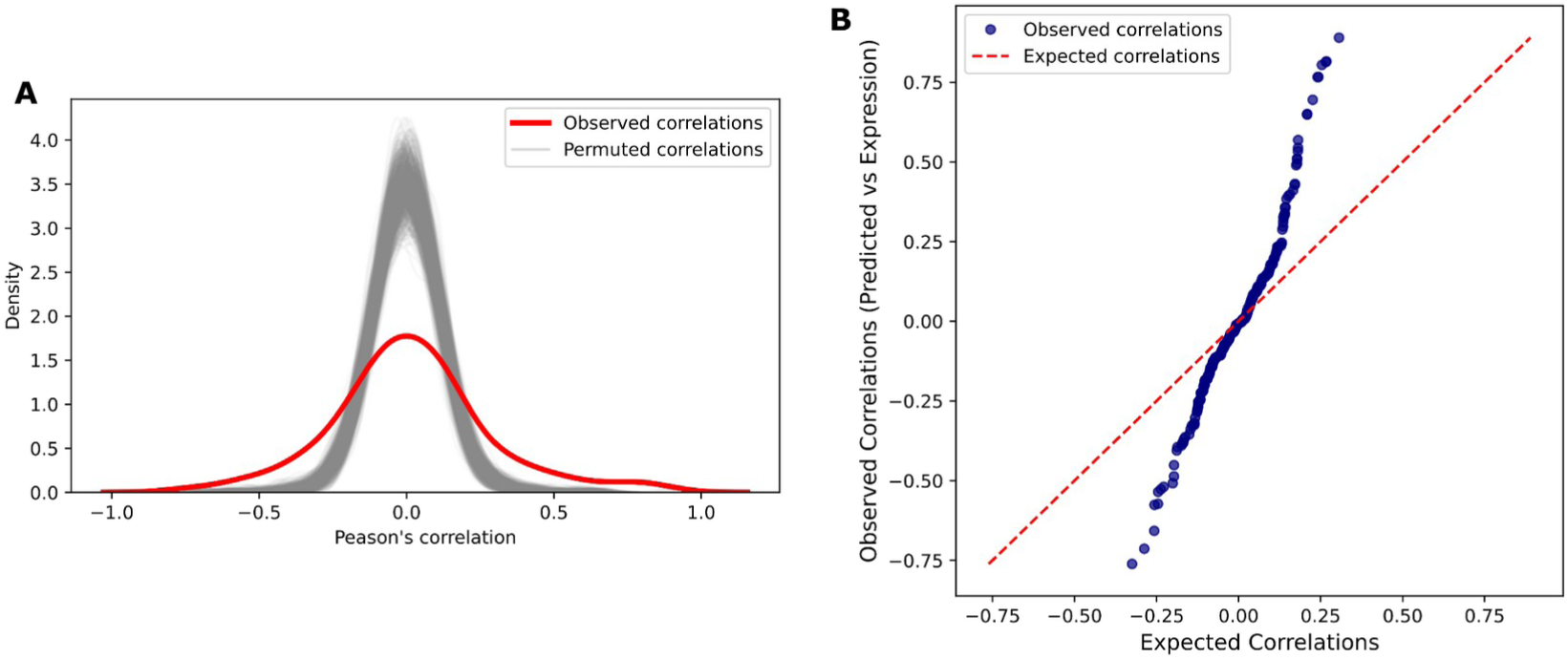
Predicted transcription rates correlate with measured gene expression in *E. coli* isolates. **(A)** The distribution of observed correlation coefficients (red curve), alongside correlation distribution from individual permutation runs (grey curves), which represent the null distribution generated by shuffling predicted transcription rates for each gene. (**B)** Represents a Q-Q plot comparing observed correlation coefficients (y-axis) to the expected correlation coefficients derived from the permutation tests (x-axis).

While the majority of genes exhibited weak correlations, 17% (N=46) of DEGs demonstrated moderate to strong positive correlations, supporting the hypothesis that these variants can regulate gene expression through their impact on promoters as observed in genes like *yfgJ* (putative oxidoreductase), *kgtP* (α-ketoglutarate transporter)*, waaQ* (heptosyltransferase involved in LPS biosynthesis), and *ulaE* (L-ascorbate catabolism enzyme)^24^ (**Figure 4**). The remaining unexplained DEGs suggest the presence of additional regulatory mechanisms, either acting in *cis*, or in *trans*.

**Figure 4:**
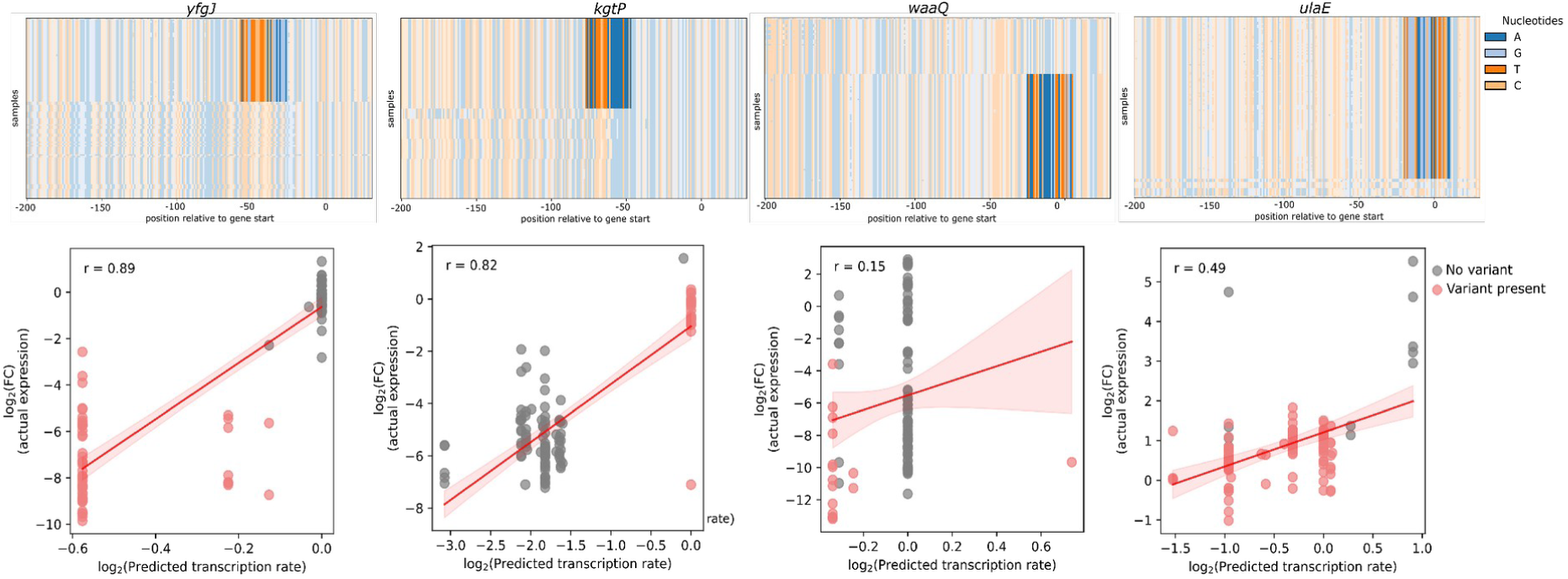
Non coding variants within canonical promoters influence predicted transcription rates and correlate with gene expression. The top panel shows the variant map for four genes (*yfgJ, kgtP, waaQ, ulaE*), with nucleotides aligned relative to the gene’s start codon. Solid color regions indicate the non-coding variant altering the predicted transcription rates. Bottom panels show the correlation between the gene’s predicted transcription rates and their measured gene expression (log_2_ fold change). Isolates with non-coding variants are highlighted in red, and those without the variant are in grey.

Similarly to canonical promoters, transcription factor binding sites (TFBSs) are important sequence-based elements that influence gene expression by allowing for the binding of specific transcription factors (TFs) that can selectively recognize their cognate TFBS and either support (activators) or repress (repressors) transcription. We compiled a list of 93 experimentally validated TFBS motifs from RegulonDB^25^ and scanned the non-coding sequences of the DEGs in each *E. coli* isolate to identify the presence or absence of each TF’s binding site. We identified the presence of at least one TFBS for 99% of the DEGs (**Supplementary** Figure 2). We then constructed a binary matrix of TFBS presence or absence across isolates and performed a GWAS to evaluate associations between TFBS variation and DEG expression.

We observed significant associations between TFBSs presence and expression for 25% of the tested DEGs (178), of which 129 (72%) had mutated σ^70^ promoters previously identified in our earlier analysis as linked to gene expression. While the presence of TFBS may suggest a potential influence on gene expression, it is important to note that the regulatory effect depends on whether the TF is an activator or repressor. Hence, our observations do not directly correlate with the upregulation or downregulation of the genes. Notably, 5% (23) of the identified TFBSs overlapped with SNP positions previously associated with gene expression. This overlap suggests that these SNPs could potentially alter TF binding by disrupting existing or creating novel TFBS, which in turn may affect gene transcription and expression, as seen in genes such as *yihM*, *hipB* (antitoxin component of a type II toxin-antitoxin system)^24^ (**Figure 5**). Through this second in-silico validation we have identified an additional mechanism driving changes in gene expression through *cis* non-coding genetic variation.

**Figure 5:**
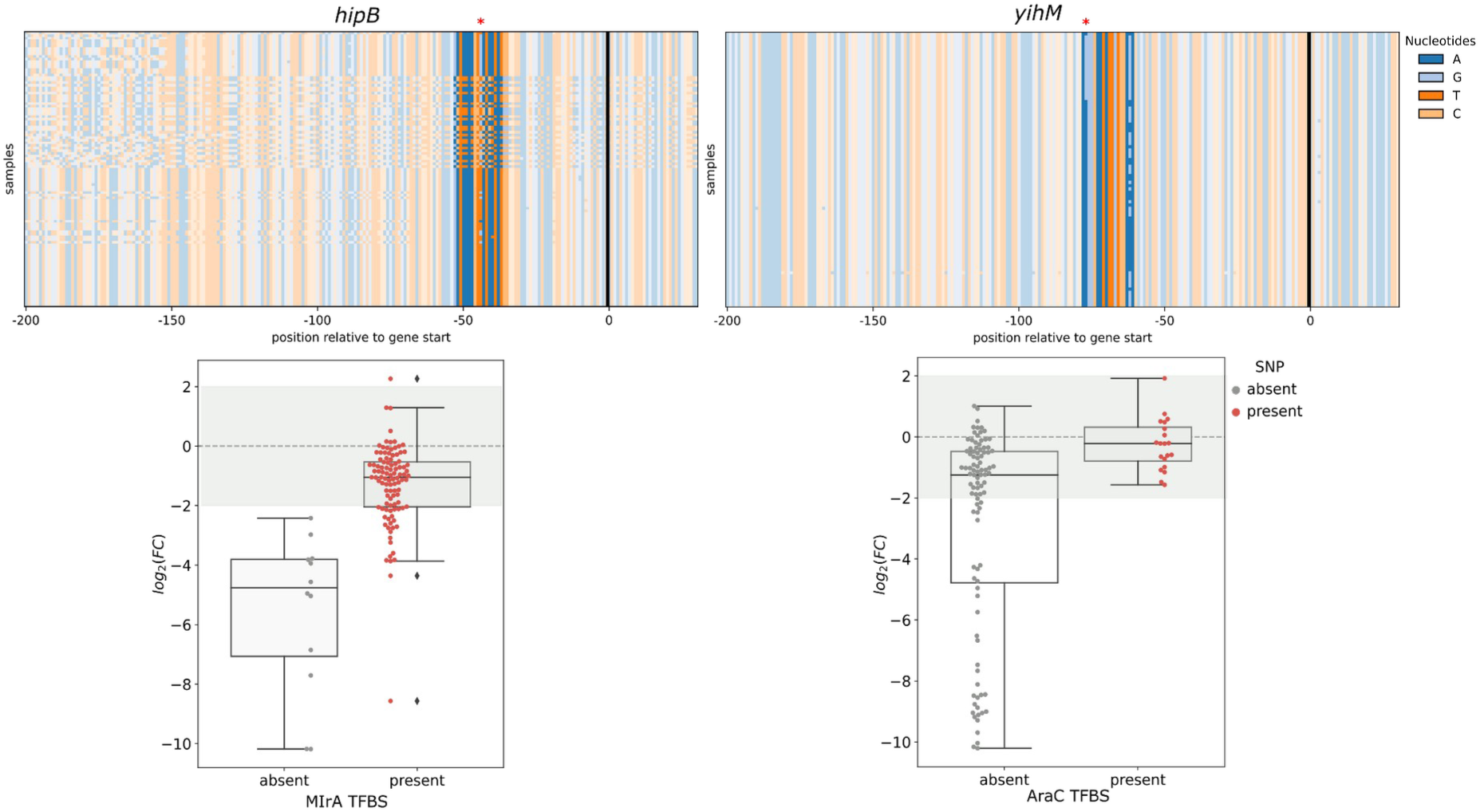
TFBS overlap with SNP positions are significantly associated with altered gene expression in *E. coli*. The top panel represents the variant maps upstream of *hipB* and *yihM* genes, aligned relative to the start codon of the corresponding gene. The red asterisks (*) indicate SNP positions previously linked to gene expression changes and overlap with the TFBS MlrA (for *hipB*) and AraC (for *yihM*). The box plots show the relationship between the SNP presence within the TFBS and gene expression. The y-axis represents log₂ fold changes in expression, while the x-axis indicates the presence or absence of the TFBS. The overlaid swarm plots show the presence (red) or absence (grey) of the variant.

We further validated a selection of the original *E. coli* associations using a fluorescent reporter assay (N=18 genes). We amplified the non-coding regions of these 18 DEGs with significant associations from the reference and from a selected isolate carrying the associated variant, and cloned them upstream of the reporter gene (GFP) into the pOT2 plasmid. Differences in the expression of the GFP gene due to the impact of the variants in gene expression would then translate to differences in observed fluorescence. The generated constructs were cultured in supplemented M9 media, and GFP expression was measured over 16 hours at 30-minute intervals. We observed a clear difference between the GFP expression of the mutated reporter constructs versus those of their wild-type counterparts for 16 genes (**Figure 6A**). Expression changes ranged from −1 to +3-fold, with effect sizes ranging from small (Cohen’s *d* ≤ 0.2, 2 genes), medium (0.2 < *d* < 0.8, 4 genes), and large effects (*d* ≥ 0.8, 12 genes). While the direction of gene expression changes between the reporter fusion assay and the actual gene expression was not always consistent with the original variation measured through RNA-seq, 9 out of 18 genes (50%) showed concordant directionality, which further validates the use of GWAS to find the genetic determinants of gene expression changes.

**Figure 6:**
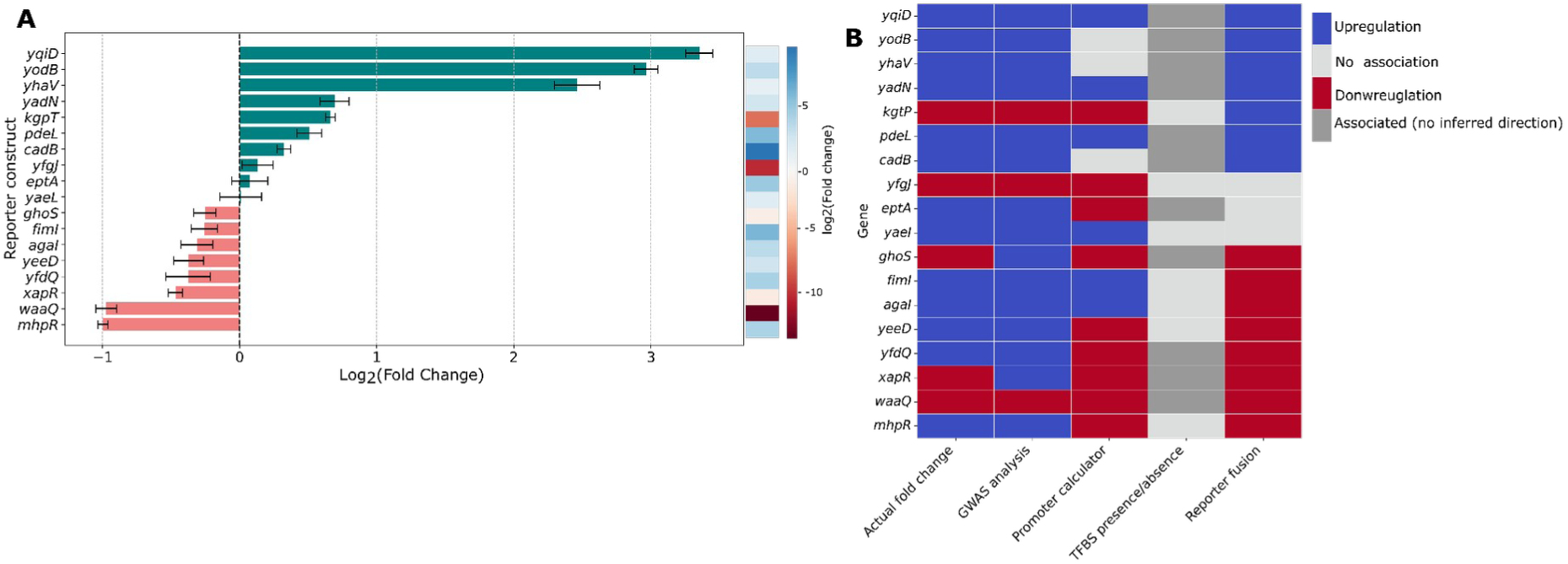
Non-coding variants alter transcription activity in *E. coli*. Reporter constructs containing either wild-type or mutated non-coding regions were generated for 18 genes. Cultures were grown for 16 hours, with GFP expression measured every 30 minutes. Fluorescence readings were normalized to both blank controls and cell density. To reduce noise observed in early measurements, only data collected after 6 hours were included in the analysis. **(A)** Relative fold-changes in GFP expression of the mutated reporter construct relative to its wild-type counterpart. For each pair, mean fluorescence values were averaged across replicates from the 6-hour time point onwards. Fold enrichment was computed as the ratio of the signal in the mutant to the one in the wild-type. The heatmap on the right shows the fold changes of DEG gene in the initial screen (rich media, exponential growth phase). **(B)** Agreement between the approaches used to establish links between non-coding variants and gene expression changes. The different color indicates the direction of gene expression change associated with each approach. For TFBSs, dark gray indicates a significant association between their presence and gene expression. The direction of the effect cannot be determined, as transcription factors may function as either activators or repressors, and this context-specific role could not reliably be inferred here.

We next compared the results from the different approaches used to establish the relationship between non-coding variants and gene expression in *E. coli*: GWAS using k-mers, canonical promoter strength predictions, TFBS presence/absence analysis, and reporter fusion assays (**Figure 6B**). For each method we categorized the predicted effect as upregulation, downregulation, or no significant change. In the GWAS analysis, beta values were used to infer directionality, in the promoter calculator, this was based on the sign of the correlation coefficient between the predicted transcription rates and measured gene expression, and in the reporter assays directionality was inferred based on log_2_ fold-change and effect sizes (upregulation with log2 fold-change > 0 and *d* ≥ 0.5, downregulation with log_2_ fold-change < 0 and *d* ≥ 0.5, or no significant change with *d* < 0.5). For TFBS presence/absence however directionality is inherently ambiguous, as the presence of a binding site does not indicate upregulation, nor does its absence imply downregulation. In this context, TFBS associations are best interpreted as indicators of regulatory involvement rather than direct predictors of the direction of expression change. Overall, 22% (N=4, *yqiD*, *pdeL*, *waaQ*, and *yadN*) of the genes showed consistency in their effects on gene expression across all three methods. However, we also observed discrepancies, such as in the case of *ghoS*, where GWAS found an association with upregulation, while the promoter calculator and reporter fusion assays predicted downregulation. Although the TFBS association aligns with higher expression, the unknown role of the TF (activator or repressor) limits functional interpretation (**Figure 6B**). The GWAS analysis showed the highest concordance with the actual gene expression changes, agreeing with observed changes in 89% of cases, which can be expected as the method uses the actual gene expression to find associations, followed by the promoter calculator which was in agreement in 61% of the cases. The reporter fusion assay and TFBS presence or absence analysis aligned with observed gene changes in 50% of cases. This variability in concordance across genes highlights the multifaceted nature of regulatory control and suggests that no single approach fully captures the complexity of expression variation driven by non-coding genetic variations.

### Antimicrobial resistance in *E. coli* and *P. aeruginosa* can be linked to genetic variations within non-coding regions

Antimicrobial resistance (AMR) poses a significant threat to global public health and is driven largely by the genetic adaptability of bacterial pathogens. While the role of protein-coding genes in resistance has been well studied, the contribution of non-coding regulatory regions remains underexplored. We therefore analyzed the genomes of *E. coli* (N=1,675) and *P. aeruginosa* (N=413, the same isolates for which we had available transcriptomic data) isolates, obtained from existing studies^10,26^, to understand to what extent non-coding variants can explain antimicrobial resistance in these pathogens.

The *E. coli* isolates were tested for susceptibility against 12 antibiotics including amoxicillin-clavulanic acid (AMC), ampicillin (AMP), cefuroxime (CXM), cefotaxime (CTX), cephalothin (CET), ceftazidime (CAZ), gentamicin (GEN), tobramycin (TOB), ciprofloxacin (CIP), amoxicillin (AMX), trimethoprim (TMP), and piperacillin-tazobactam (TZP). Resistance profiles varied significantly, with AMP showing the highest resistance rate (75% of isolates resistant), while TZP exhibited the highest susceptibility rate (60% of the isolates susceptible) (**Supplementary** Fig 3A). For *P. aeruginosa*, among the four antibiotics, CIP, TOB, CAZ, and meropenem (MEM), tested. TOB showed the highest sensitivity rate (67%) and MEM showed the highest proportion of resistant isolates (74%) (**Supplementary** Figure 3B).

We identified SNPs and InDels within the non-coding regions of these isolates, as previously stated, and conducted a whole-genome GWAS (wGWAS) to identify resistance-associated variants across the genome as well as a *cis*-GWAS focusing only on non-coding variants near genes. Both approaches identified resistance-associated variants in both *E. coli* and *P. aeruginosa* (**Supplementary** Figure 4 to 9).

In *E. coli* the wGWAS and *cis*-GWAS identified on average 137 and 44 genes, respectively, encoding resistance-associated variants across all antibiotics except for amoxicillin (AMX) (**Figure 7A**). The *cis* non-coding variants were found in poorly characterized genes like *yghJ*, *yieH*, and *osmC*, which are implicated in biological processes such as lipoprotein secretion and stress responses and have been previously recognized as markers for predicting AMR in *E. coli*^24^. We also observed that some of the *cis* non-coding variants were similar across different antibiotic classes with the most overlap being between cephalosporins and aminoglycosides (**Figure 7C**). Some variants appeared to play broader roles in resistance across multiple drug classes. For example, non-coding variants in *gadX* were associated with resistance to cephalosporins, aminoglycosides, and fluoroquinolones. GadX is a transcriptional regulator known to enhance expression of the MdtEF efflux pump, which contributes to multidrug resistance^27^. Similarly, non-coding variants in *mbsA*, a homolog of the multidrug resistance ATP-binding cassette (ABC) transporter, were identified in isolates resistant to both cephalosporins and aminoglycosides^28^. This cross-class overlap of non-coding variants suggests their potential role in mediating cross-resistance mechanisms among different antibiotic classes.

**Figure 7:**
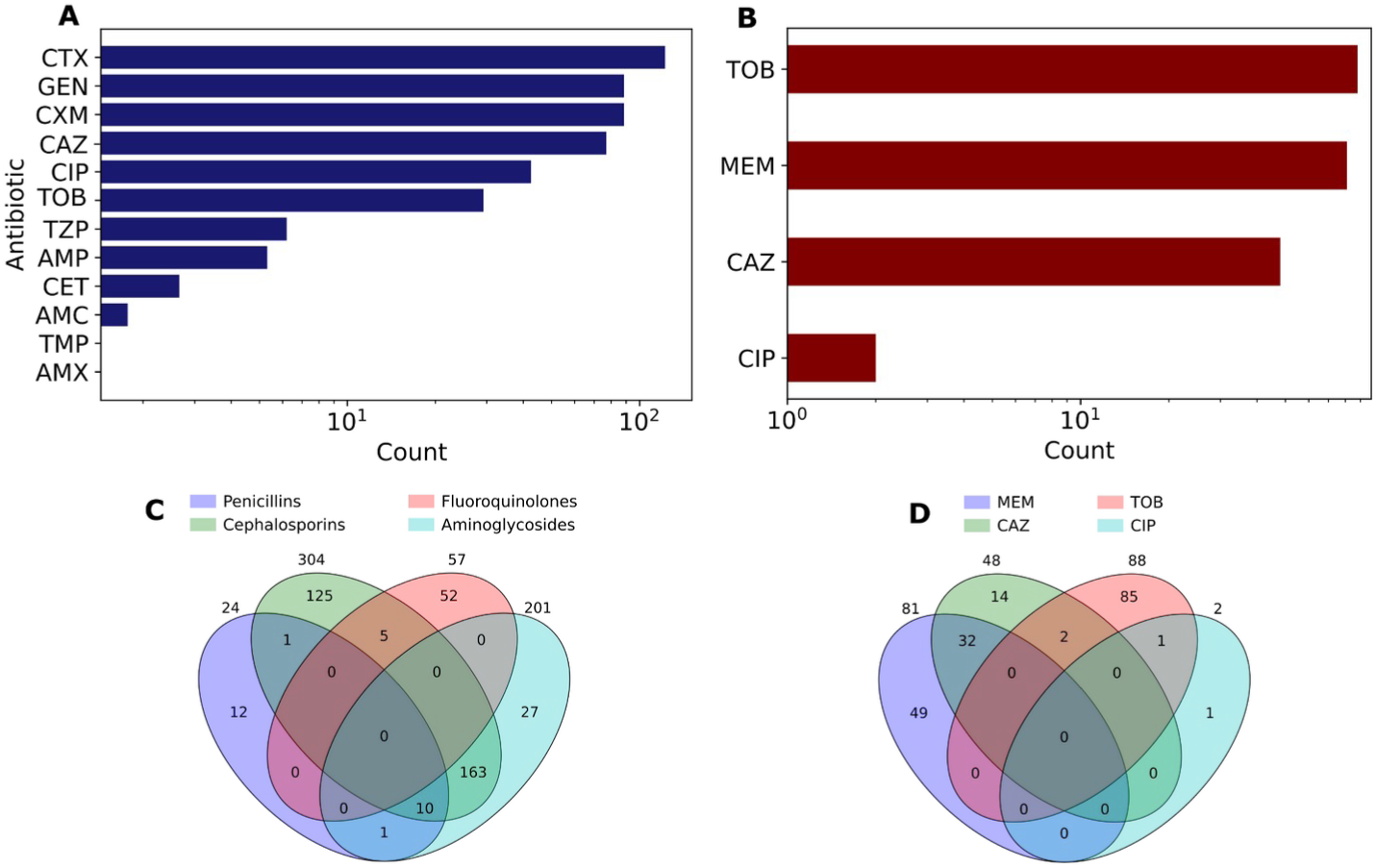
Non-coding variants are associated with antimicrobial resistance in *E. coli* and *P. aeruginosa*. Panels **A)** and **B)** show genes with non-coding variants associated with antibiotic resistance. The y-axis represents the antibiotic and the x-axis represents the number of genes with non-coding variants associated with resistance. **C)** and **D)** represent Venn diagrams that show the overlap and uniqueness of non-coding variants associated with antibiotic resistance across different antibiotics.

In *P. aeruginosa*, we identified on average, 93 genes and 27 genes, for wGWAS and *cis*-GWAS respectively, with resistance-associated variants across all four antibiotics. Among the antibiotics tested, we observed the highest number of genes with resistance-associated non-coding variants for TOB (**Figure 7B**). Some of these non-coding variants were located in the *rpSA* gene which has been linked to translation initiation in *E. coli* and pyrazinamide resistance in *Mycobacterium tuberculosis*^29–31^. However, its specific role in mediating resistance in *P. aeruginosa* remains unclear. We also identified a non-coding variant in the *murU* gene, which encodes N-acetylmuramate α-1-phosphate uridylyltransferase, which is involved in peptidoglycan recycling and has been linked to antibiotic resistance in *P. aeruginosa* through its regulation of cell wall precursor pools^32^. Some of the non-coding variants were shared between CAZ and MEM as well as between TOB and CAZ, suggesting a potential cross-resistance mechanism between these antibiotic classes (**Figure 7D**). This cross resistance could result from overlapping regulatory pathways that influence resistance to these different antibiotics.

### Non-coding variants confer tobramycin resistance by affecting bacterial cell wall and outer membrane composition

We identified a SNP in *E. coli*, a T>C transition, at position 3808241 upstream of the *waaQ* gene, which is part of the *waa* operon. This operon includes nine other genes, *waaG*, *waaP*, *waaS*, *waaB*, *waaO*, *waaJ*, *waaY*, *waaZ*, and *waaU*, which collectively contribute to lipopolysaccharide (LPS) synthesis^24^. The *waaQ* gene encodes a heptosyltransferase responsible for transferring heptose molecule (III) to the inner core of LPS, which is a critical step in its biosynthesis^33,34^.

We selected three strains harboring the SNP and three strains without the variant, including the *E. coli* K-12 reference strain. Strains bearing this variant exhibited higher MIC values for TOB (1-8 µg/mL) than those without the variant (0.125–0.5 µg/mL) (**Supplementary Table 8**). To understand the impact of this variant on *waaQ* expression under antibiotic stress, we exposed the strains to ½ MIC for TOB, a sub-inhibitory concentration that is sufficient to induce physiological responses without completely halting bacterial growth. Differential expression analysis of the *waa* operon genes revealed distinct patterns between strains with and without the variant. Two variant-bearing strains (IAI76 and IAI78) showed downregulation of the *waa* operon, whereas one strain (IAI77) displayed incomplete expression of four out of ten *waa* genes (**Figure 8A**). This suggests that the non-coding SNP may influence not only *waaQ* expression, but also the coordinated regulation of other genes in the operon. Gene interaction analysis using the STRING database^35^ identified significant links between the *waa* operon genes and known AMR genes *ugd*, and ompF (**Figure 8B**). The *ugd* gene has been tied to LPS modification, resulting in the reduction of antibiotic binding, and uptake^28^. OmpF is a porin that facilitates the diffusion of small hydrophilic molecules across the bacterial membrane, and is frequently associated with reduced antibiotic uptake owing to mutations, as well as being downregulated upon the deletion of *waaF*^28,36,37^. Notably, *ompF* was upregulated in one strain (IAI76), which also exhibited the most pronounced downregulation of the *waa* operon genes. These findings highlight the potential connection between the *waa* operon and AMR pathways involving LPS modification and membrane permeability.

**Figure 8:**
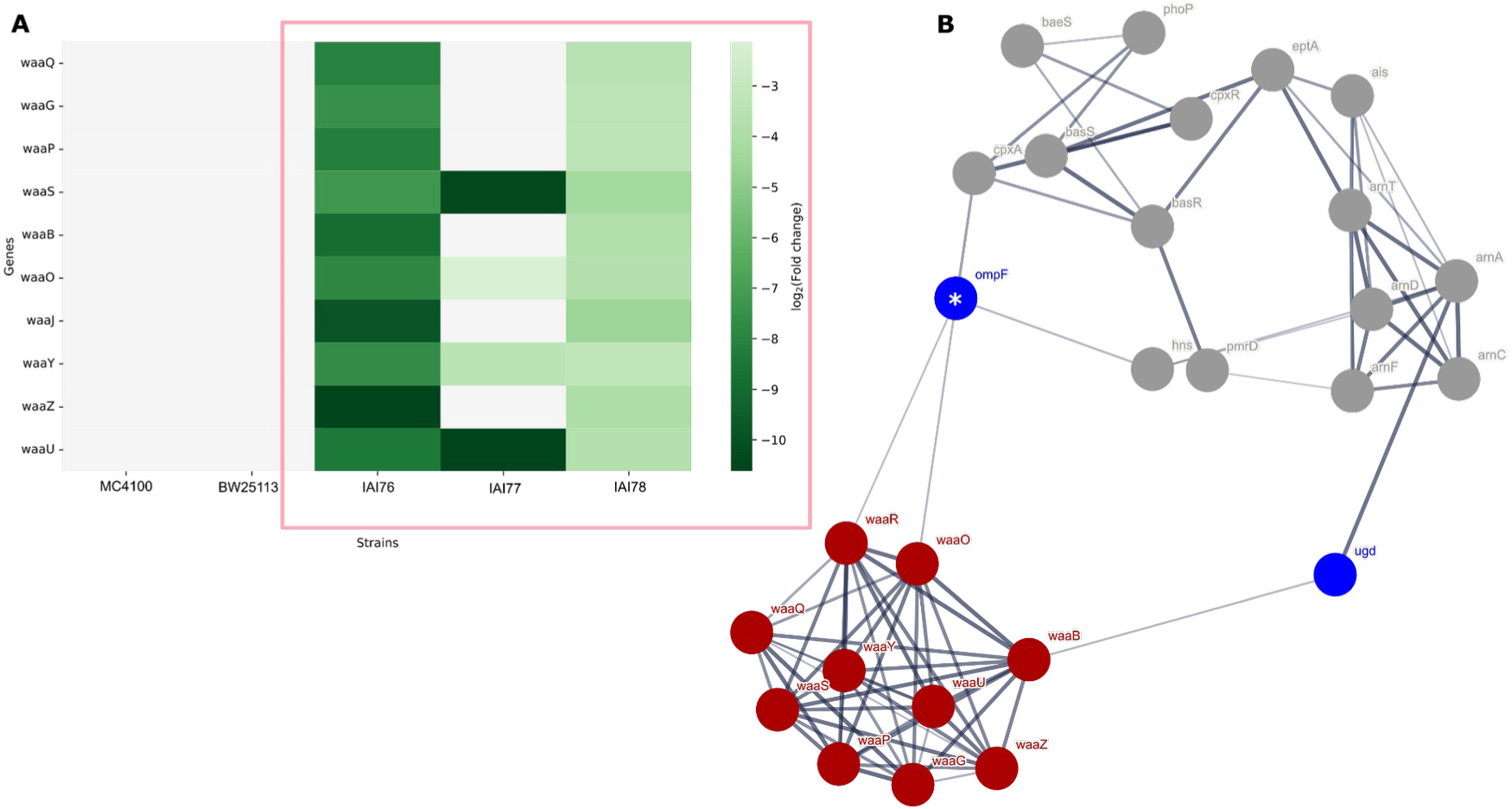
The expression of *waa* operon genes varies per strain with the operon showing connections with known antimicrobial resistant genes in *E. coli*. Six strains (three strains bearing the *waaQ*/3,808,241/T>C variant and three without) were exposed to ½ MIC for TOB for one hour. RNA was extracted after the exposure and sequenced. Transcriptomic analysis was conducted by comparing the expression of genes in each strain with that of the reference strain. **(A)** Expression of the *waa* operon across control and test (red rectangle) strains. A gene was indicated as differentially expressed if it had a log2fold change ≤ −2 or ≥ 2 and an adjusted p-value < 0.0001. **(B)** Interaction between *waa* operon genes (in red) and known antimicrobial-resistant genes (ARGs) (in blue). The asterisk (*) denotes ARGs that are differentially expressed in at least one strain.

We also investigated the SNP, G>T transversion, at position 670,467, upstream of the *murU* gene in *P. aeruginosa*, which was associated with increased resistance to TOB. Six strains were exposed to ½ MIC of TOB for one hour. Strains harboring the *murU* non-coding variant (n=3) exhibited significantly higher TOB resistance (>16 µg/mL) compared to those without this variant (n=3, including the reference strain PA14) (**Supplementary Table 8**). From our differential expression analysis, we did not observe significant changes in *murU* expression between the variant and non-variant strains following TOB exposure.

## Discussion

The ability to use biological sequences alone to determine the characteristics of a living organism has long inspired researchers to identify general rules that would allow for the prediction of phenotypes from primary sequence alone. Fundamentally, this problem can be tackled using two complementary approaches. Mechanistic approaches require a certain level of understanding of the functioning of molecular systems, sometimes down to the level of individual molecules, making its application beyond individual genes or model species prohibitive. In contrast, statistical approaches on the other hand do not require an “intimate” understanding of the system under study, and thus can be applied to a wide variety of molecular processes and species. They do however require a higher upfront “cost” via the accumulation of sufficient observations to achieve an appropriate level of statistical power.

Because of the high coding density of bacterial genomes, the vast majority of genotype-to-phenotype research in this kingdom has focused on predicting the consequences of changes to coding sequences^38^. However, variation in gene expression is likely to be as important in explaining phenotypic changes in bacteria, as exemplified in numerous studies concerning basic physiology^39,40^, as well as phenotypes related to infection research^41^, such as AMR^42^ and pathogenicity^11^. Identifying the extent to which non-coding variants influence gene expression variation could then improve the prediction of phenotype from primary sequence, with positive consequences for diagnostics and synthetic biology.

In this study we have focused on genetic variants in *cis* non-coding regions, under the assumption that in bacterial species they would be responsible for most of the variability in gene expression. We encoded these variants using three distinct data structures, thus ensuring that we would be able to account for all kinds of genetic variations. The use of k-mers in particular, allowed us to jointly consider short variants, gene and promoter presence/absence patterns, and promoter switching, a phenomenon that is estimated to affect up to 16% of DEGs across *E. coli* isolates^9^. We then applied statistical genetics to identify associations between variants and gene expression changes. In the case of *E. coli*, we used the accumulated knowledge on σ^70^ and TFBS sequences to validate the associations mechanistically, as well as experimentally for a smaller subset.

Consistent with the differences in the genetic structure of the two species, we found differences in the proportion of DEGs, for which we identified a variant associated with expression changes. These differences dissipated when using k-mers, for which we found associations for 39% of DEGs in both species, and were only apparent when using short variants and especially promoter switching events. For the latter variant category we could identify associations only for 2% of DEGs in *P. aeruginosa*, as opposed to 28% in *E. coli*. We found the opposite picture when looking at short variants, with associations for 34% of *P. aeruginosa* DEGs, and 7% only for *E. coli*. From these observations we can conclude two things. The first, methodological in nature, confirms that encoding genetic variants as k-mers allows to capture the most genetic variation; the second is that different species have different sources of genetic variation fueling adaptation.

Using external^26^ (*E. coli*) and overlapping^10^ (*P. aeruginosa*) datasets on susceptibility to antimicrobials, we have shown that non-coding variation is also associated with downstream phenotypes with high clinical relevance. In both species, about one third of the genes with genetic variants (in any region of the genome) associated with drug susceptibility, had these variants encoded in non-coding regions. In *E. coli*, we identified resistance-associated non-coding variants in both known AMR genes such as *gadX*^27^ and in genes not previously linked to resistance, such as *waaQ*, a component of the *waa* operon involved in LPS synthesis^24^. Strains with a T>C transition at position 3,808,241 upstream of *waaQ* showed downregulation of the *waa* operon when exposed to sub-inhibitory concentrations of tobramycin. Given the role of LPS in membrane permeability and the observed protein interactions with resistance-related genes *ompF* and *ugd*^28,36^, we suggest that this variant may contribute to tobramycin resistance through modification of the outer membrane. For *P. aeruginosa*, we specifically tested the effect of G>T transversion at position 670467, a non-coding variant found in *murU*, a gene involved in peptidoglycan recycling, a mechanism shown to contribute to resistance in the organism^32^. Although we did not observe differential expression of the *murU* gene in variant-bearing strains, the absence of an effect may reflect context-dependent compensatory regulation provided by the strain background or insufficient antibiotic exposure.

Even though we have identified a large number of non-coding genetic variants associated with gene expression and AMR, our study has a number of limitations. The limited availability of transcriptomic data reduced the statistical power of our association analyses, especially in *E. coli*. Future studies with larger datasets could enhance the robustness of our findings, uncover rarer variants, and associations with more subtle regulatory effects that may have been masked in our study. While collating data from public databases such as ENA and GEO may be a cost-effective way to increase sample size, questions on the homogeneity of experimental and sequencing protocols make this exercise not free from bias. The application of advanced machine learning such as convolutional neural networks^43^ (CNNs) and foundation models^44^, both able to operate on primary sequence, also holds promise for the identification of genetic variants influencing gene expression. Both would require large datasets for training or even just for validation. Lastly, the lack of well-characterized transcriptional regulatory elements for *P. aeruginosa* prevented the *in-silico* validation of associations in this species. Even for *E. coli*, for which we could make use of the decades of work on characterizing its regulatory network, we could not always validate the associations we identified, which could imply that variants outside of *cis* non-coding regions have a sizable impact on gene expression. Providing a mechanistic model for how those trans variants affect gene regulation will likely be even more difficult.

## Methods

### Bacterial strains

Whole genome sequences from 117 *E. coli* isolates, and 413 *P. aeruginosa* isolates were obtained from two studies^10,15^. RNA-seq data for *P. aeruginosa* was also obtained from the study from which we obtained the genomes, but for *E. coli*, the data was generated as part of this study. RNA was extracted from exponentially growing bacteria in rich media (LB), with samples being harvested at an OD_600_ of 0.2 (**Supplementary Table 4**). These datasets were analyzed to decipher the relationship between non-coding genetic variants and gene expression in these isolates. For *P. aeruginosa*, data on the susceptibility of each isolate were available for four antibiotics: ciprofloxacin (CIP), gentamicin (GEN), meropenem (MEM), and tobramycin (TOB). We also included an independent dataset^26^ of 1,675 *E. coli* isolates together with their susceptibility profiles for 12 antibiotics: Ampicillin (AMP), Cefuroxime (CXM), Cefotaxime (CTX), Cephalothin (CET), Ceftazidime (CTZ), Gentamicin (GEN), Tobramycin (TOB), Ciprofloxacin (CIP), Amoxicillin-clavulanate (AMC), Amoxicillin (AMX), Tazobactam (TZP) and Trimethoprim (TMP). These datasets were used to establish a relationship between non-coding variants and antimicrobial resistance phenotype.

### Differential gene expression and functional enrichment analysis

The raw RNA-Seq reads were first assessed for quality using FastQC v0.11.9^45^ and aligned to their respective reference genomes (*E. coli* K-12 substr. MG1655, RefSeq accession: NC_000913.3; *P. aeruginosa* UCBPP-PA14, RefSeq accession: NC_008463.1). Transcript quantification was performed using kallisto v0.46.2^46^. Differential gene expression analysis was conducted using DEseq2 v1.44.0^47^ for *E. coli* isolates and LPEseq v0.99.5^48^ for *P. aeruginosa* isolates, since a single replicate was available for this species. Significance thresholds to identify differentially expressed genes were set as follows: genes needed to be present in at least 10% of isolates to be considered, an adjusted p-value < 0.0001, and an absolute log_2_fold change ≥ 2 in at least 5% of the strains. A cluster of orthologous groups (COG) enrichment analysis was performed to highlight the main biological processes influenced by DEGs. Significance was set at an adjusted p-value < 0.05.

### Characterization of genetic variants within non-coding regions

Three different genetic markers were used to identify and characterize non-coding genetic variants: short variants (such as SNPs, single nucleotide polymorphisms, InDels, short insertions or deletions), gene cluster specific k-mers, and the presence/absence of intergenic regions (IGRs). For short variants, we performed a whole-genome variant calling analysis using snippy v4.6.0^49^ with default parameters, using the reference genome of each species for mapping. We defined a non-coding region as a 230 bp sequence spanning 200 bp upstream from the start codon and 30 bp downstream. Bedtools v2.30.0^50^ was used to select short variants occurring only within this 230 bp window. BCFtools v1.13^51^ was then used to merge the non-coding variants for each gene across the isolates, which were used as inputs for the association analyses.

A pangenome analysis was conducted for the isolates of both bacterial species using panaroo v1.5.0^52^ with default parameters, except for the following arguments: “--clean-mode strict” and “--remove-invalid-genes”. We used piggy v1.5^53^ to perform a comparative analysis of IGRs across the bacterial isolates. The gene presence/absence matrix generated by panaroo was used as input, and the analysis was performed using default parameters, except for the change in the percentage length identity to 10. We further used the gene presence/absence matrix generated by panaroo to extract gene cluster specific k-mers from the genomic regions flanking each differentially expressed gene using panfeed v1.6.1^14^. The analysis was conducted using the following parameters: “--upstream 200”, “--downstream 30”, “--no-filter”, “--maf 0.1”, “--downstream-start-codon”, and “--consider missing”. The resulting presence/absence from piggy and kmer unique patterns were used as inputs for the association analyses.

### Phylogenetic analysis

The phylogenetic tree of the strains was computed using ParSNP v1.2^54^, which generated a whole genome nucleotide alignment for all the strains. The resulting tree was visualized using iTOL v6^55^.

### Associating non-coding variants with gene expression and resistance phenotypes

We performed a GWAS analysis using pyseer v1.3.9^56^, to understand the relationship between non-coding genetic variants and gene expression, and antimicrobial resistance phenotypes. For the gene expression phenotype, we examined associations between the presence or absence of non-coding genetic variants (captured by each set of genetic markers) and differentially expressed genes. Variants with minor allele frequencies (MAF) below 10% were excluded from this analysis. Linear mixed models, as implemented in pyseer, were employed to account for genetic relatedness among the isolates, ensuring the robust detection of associations. The population structure was corrected using a similarity matrix generated from the generated phylogenetic tree. To address the issue of multiple testing inherent in GWAS analysis, we applied a pattern-based method that adjusted for the number of unique tests performed. This reduces the risk of over-correcting the association p-values, as it is based on the number of unique tests being carried out. In linking these variants with resistance phenotype, we considered all genes within the bacterial genome, regardless of whether they were differentially expressed. For *E. coli* used a binary phenotype (1-resistance, 0-susceptibility) as the previous authors did, and for *P. aeruginosa* we made use of (MIC) values. This GWAS was conducted in a similar manner as previously described, but with a lower MAF threshold of 0.01 to include rarer variants.

### Validating associations between non-coding variants and gene expression in *E. coli*

#### In-silico validation

We used a modified version of the Promoter Calculator^13^, a biophysical model developed to estimate σ^70^ dependent transcription rates in *E. coli*. We focused on genes where k mers were associated with gene expression, under the hypothesis that these variants might influence transcription by altering the canonical σ⁷⁰ promoter sequence. The predictions were made on 230 bp sequences across the strains. The predicted rates were normalised for each gene by dividing the value for each isolate by that of the reference strain, followed by a log _2_ transformation. A Pearson coefficient was then computed between these values and the corresponding measured gene expression levels across strains. To assess whether these patterns were stronger than expected by chance, we randomly shuffled the predicted rate values 1,000 times across isolates for each gene. A new set of Pearson correlation coefficient was computed between these values and the measured gene expression values. We opted for a global approach to capture broader trends in transcriptional regulation driven by the non-coding variation in promoter regions. This also avoided the burden of multiple testing and variability that may be introduced by per gene analyses. A Kolmogorov–Smirnov test was performed to validate if the observed distribution of correlation coefficients was significantly different from the null distribution. To further validate these associations, We retrieved 93 experimentally validated TFBS from RegulonDB^25^ and used FIMO^57^ to check for the presence of these sites within non-coding sequences. We focused on significant TFBS (adjusted p-value < 0.01, present in at least 10% of the *E. coli* strains). We generated a matrix indicating the presence (1) or absence (0) of TFBS within each gene’s non-coding region, which we used to associate with gene expression as previously stated using an MAF of 10%.

#### In vitro validation

##### Construction of reporter fusion

The pOT2 plasmid (Addgene plasmid #14460) was extracted from 16 hour cultures using a Zyppy plasmid extraction kit (Cat. No. D4019, Zymo Research) following the manufacturer’s instructions. The extracted plasmid was linearized using *HindIII* FastDigest restriction enzyme (Cat. No. FD0504, Thermo Fisher Scientific, USA), according to the manufacturer’s instructions. The linearized vector was separated from other fragments using 0.7% agarose gel and purified using the Monarch DNA Gel Extraction kit (Cat. No. T1020S; New England Biolabs, UK). The 230bp non-coding regions, as defined above, were amplified using specific primer sets under conditions specified in (**Supplementary Table 5**), and the amplicons were cleaned using Monarch PCR and DNA Clean up Kit (Cat. No. T1030S; New England Biolabs, UK). To generate reporter constructs for each gene, we set up a reaction mixture containing 2 μL of 5X In-Fusion Snap Assembly Master Mix (Cat. No. 638951, TaKaRa), 6.6 μL deionized water, and linearized vector and insert at a 1:2 ratio. The mixture was then incubated at 50°C for 15 min in a thermocycler and stored on ice. The assembled product was transformed into chemically competent *E. coli* DH5-alpha cells (Cat. No. 638951; TaKaRa) via heat shock. Transformed cells were plated on LB agar containing 10 μg/mL gentamicin and incubated overnight at 37°C. The following day, colonies were screened by colony PCR using the forward primer of the insert and reverse primer of the vector (**Supplementary Table 2**). Positive clones were sent off for whole plasmid sequencing by Eurofins Scientific to confirm successful cloning.

##### Measurement of fluorescence in reporter fusions

Each version of the reporter construct (wild-type and mutated) was streaked onto LB agar containing 10 μg/mL gentamicin and incubated overnight at 37°C. Single colonies were grown in 2 mL of freshly prepared M9 medium supplemented with 0.5% glucose, 0.1% casamino acids, 0.01 mg/mL thiamine, 0.123 mg/mL MgSO_4_, 0.0147 mg/mL CaCl_2_, 0.0028 mg/mL FeSO_4_.7H_2_O, 10 μg/mL gentamicin. The cultures were incubated overnight at 37°C with shaking at 250 rpm. The next day, cultures were diluted in supplemented M9 media to a final OD₆₀₀ of 0.01 and transferred into black-walled, clear-bottom 96-well plates for growth and fluorescence monitoring. Measurements were taken at a 30-minute interval for 16 h using a Synergy H1 microplate reader with excitation at 395 nm and emission at 509 nm, with each condition tested in triplicate. To assess the effect of non-coding variants on GFP expression, the GFP levels of the mutated reporter constructs were compared to those of their wild-type counterparts. GFP fluorescence measurements beginning at the 6-hour time point were averaged across replicates, normalized to background signal and cell density. The fold change in GFP expression was determined by comparing the mean normalized GFP values of the mutated constructs to those of the wild-type constructs. Cohen’s *d* test was applied to compute the effect size and provide the magnitude of differences between the wild-type and mutated reporter constructs.

### Antibiotic exposure and RNA extraction

The minimum inhibitory concentration (MIC) of TOB for each bacterial strain was established using the standard microdilution method. Overnight bacterial cultures were diluted at a 1:100 dilution factor in Mueller-Hinton (MH) medium and grown to an OD_600_ of 0.6 at 37°C with shaking at 250 rpm. Sub-inhibitory concentrations (½ MIC) of TOB were added to each culture and incubated for an additional hour. Cells were pelleted and treated with RNAprotect (Cat. No. 76506) and stored at −80°C for subsequent extraction using the RNeasy Plus Micro Kit (Qiagen, Cat. No. 76506), according to the manufacturer’s instructions. All treatments were performed in duplicate to ensure reproducibility. RNA sequencing was performed by Novogene using the Illumina NovaSeq X Plus platform, producing approximately 2Gb of raw paired reads per sample. The quality of the raw reads was accessed using FastQC v0.11.9^45^. The reads were aligned to their respective reference genomes, and transcript quantification and DEG analysis were conducted as described above.

### Gene interaction analysis

Protein-protein interaction analysis between *waa* operon and known antimicrobial resistant genes was performed using STRING v12.0^35^ web based software. The minimum interaction score was set to 0.4 (moderate confidence), while eliminating text mining as an interaction source. The resulting network was visualized directly through the STRING web interface.

## Data availability

The RNA-Seq data generated for the 117 *E. coli* isolates have been deposited in the European Nucleotide Archive (ENA) under accession number PRJNA520928. Transcriptomic data for *E. coli* and *P. aeruginosa* isolates exposed to sub-inhibitory concentrations of tobramycin are also available in the ENA under accession number PRJEB83662. Source code for statistical analysis and the generation of figures are available at: https://github.com/microbial-pangenomes-lab/bacterial_transcription

## Author contributions

MG conceived the study, with input from BFD, MZ generated the transcriptomic data for *E. coli* with supervision from AT. JE provided support for analyzing the *P. aeruginosa* transcriptomic data under the supervision of SH. BFD analyzed the transcriptomic and genomic data, conducted all the validation experiments, and produced all the figures. BFD and MG wrote the manuscript. All authors provided comments and edits and approved the final version of the manuscript.

## Acknowledgments

B.F.D. and M.G. were supported by the Deutsche Forschungsgemeinschaft (DFG, German Research Foundation) under Germany’s Excellence Strategy – EXC 2155 (Project No. 390874280). B.F.D. were further supported by the Hannover Biomedical Research School (HBRS) and the Centre for Infection Biology (ZIB) and by the Graduate School Scholarship Programme (GSSP) from the German Academic Exchange Service (DAAD). We would like to acknowledge Nick Goldman and Pedro Beltrao from the European Bioinformatics Institute (EMBL-EBI) for their valuable support in generating the transcriptomic data used in this study.

