## Supplementary Figures for "C*is* non-coding genetic variation drives gene expression changes in the *E. coli* and *P. aeruginosa* pangenomes"

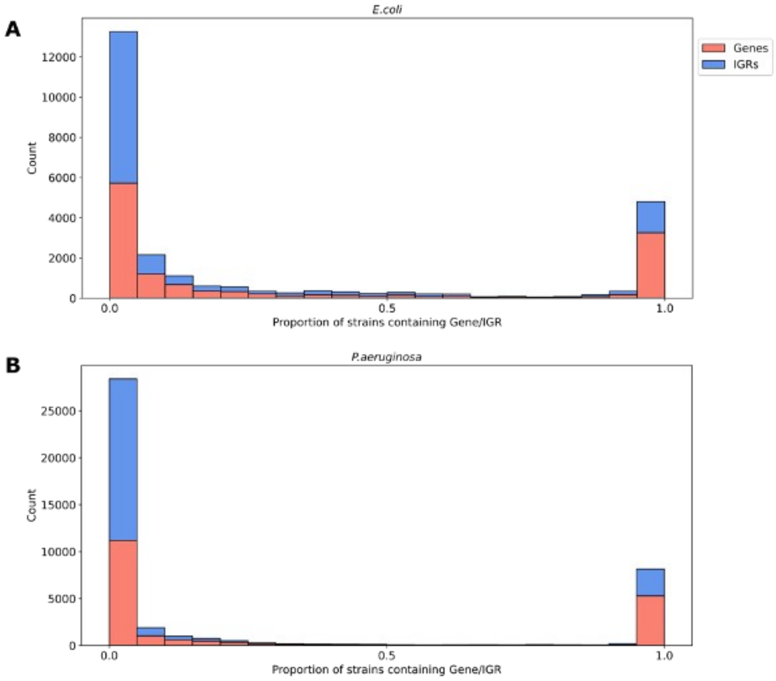


**Supplementary Figure 1: Distribution of gene and intergenic region (IGR) presence across *E. coli* and *P. aeruginosa* strains.** The number of genes (red) and intergenic regions (IGRs, blue) across isolates in *E. coli* **(A)** and *P. aeruginosa* **(B)** is shown.

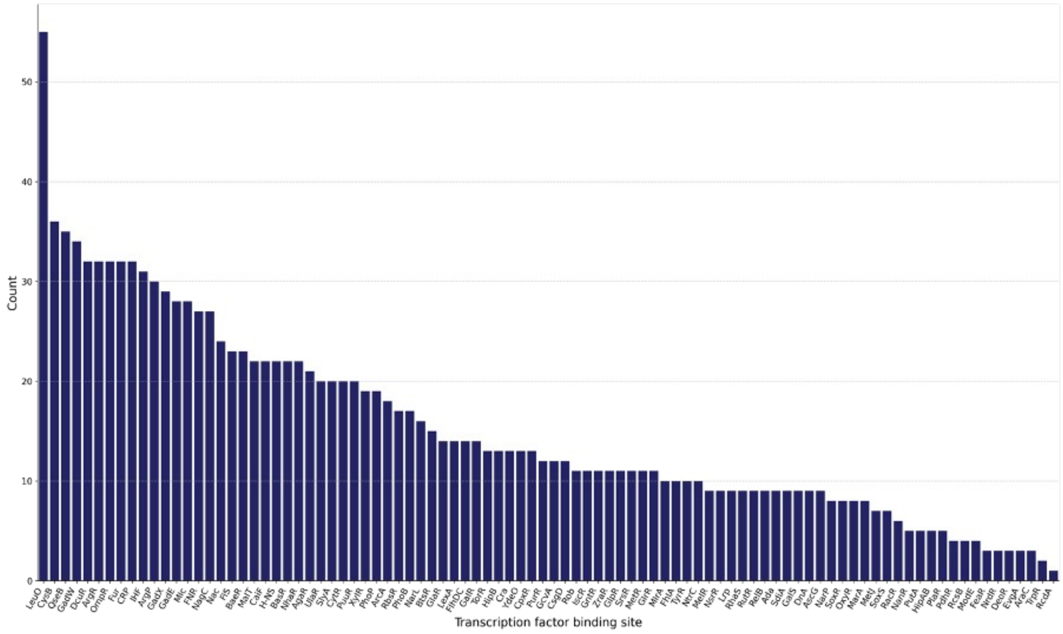


**Supplementary Figure 2: Frequency of transcription factor binding sites across non-coding regions in *E. coli* genomes.** The bar chart shows the number of differentially expressed gene where each transcription factor (TF) has a predicted binding site (TFBS) present in at least 5% of the *E. coli* isolates. TFs are ordered from most to least frequent by the number of such genes.

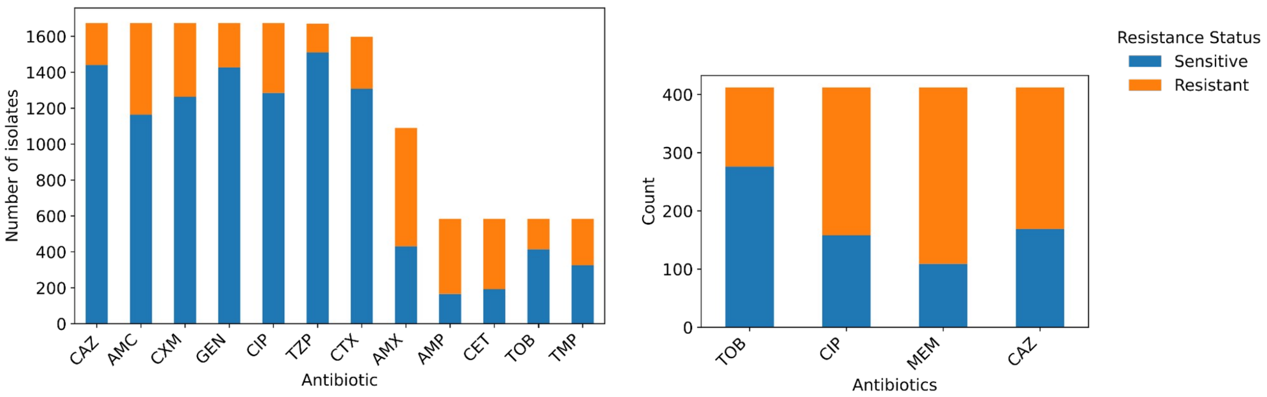


**Supplementary Figure 3: Distribution of antimicrobial resistance phenotypes for bacterial isolates**. Resistance outcomes of *E. coli* isolates to 12 antibiotics **(A)** and *P. aeruginosa* isolates to four antibiotics **(B)** represented as stacked bars indicating the number of sensitive (blue) and resistant (orange) isolates per antibiotic. The total number of isolates tested per antibiotic varies for *E. coli*, and antibiotics include CAZ (ceftazidime), AMC (amoxicillin-clavulanic acid), CXM (cefuroxime), GEN (gentamicin), CIP (ciprofloxacin), TZP (piperacillin-tazobactam), CTX (cefotaxime), AMX (amoxicillin), AMP (ampicillin), CET (ceftriaxone), TMP (trimethoprim), TOB (tobramycin), MEM (meropenem)and CAZ (ceftazidime).

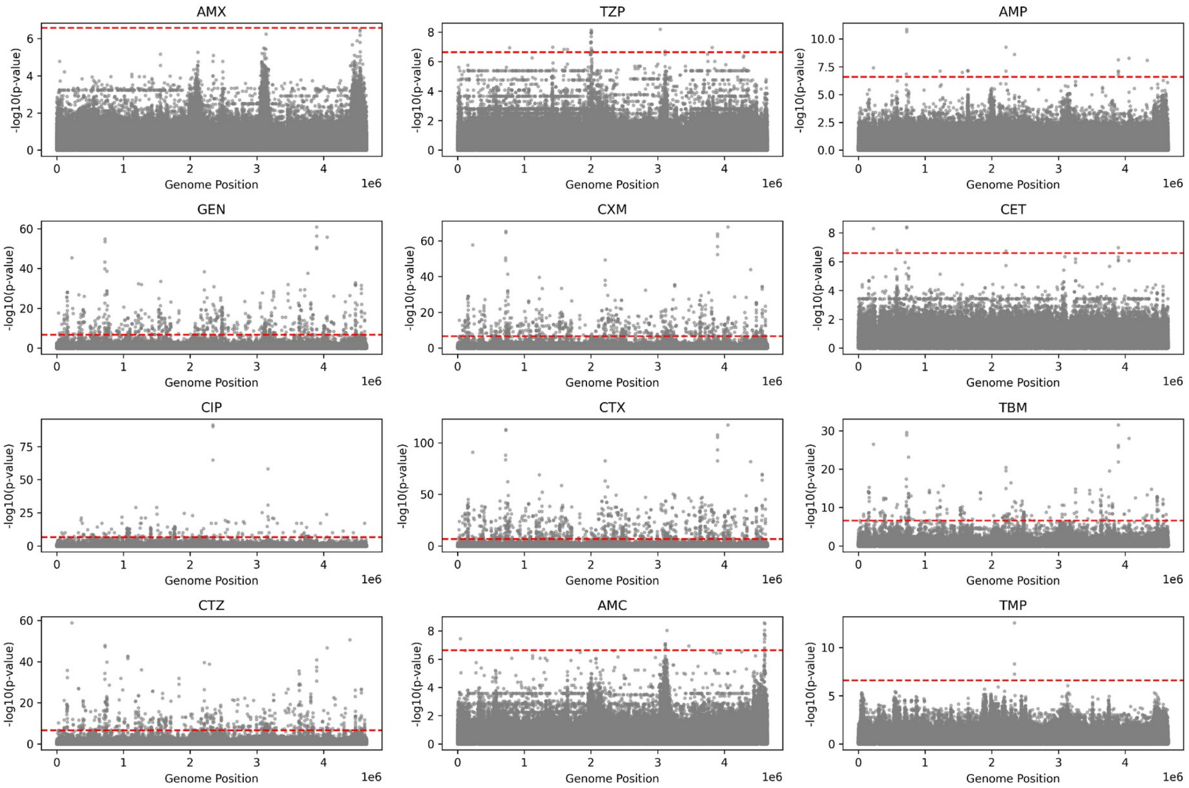


**Supplementary Figure 4: Manhattan plots for whole-genome GWAS in *E. coli.*** Each plot shows the association between an antibiotic and the resistance phenotype. Each grey dot represents a SNP, with the x-axis indicating the variant's position in the genome. The y-axis shows the p-values corrected for multiple testing. Variants above the red dashed line are considered statistically significant.

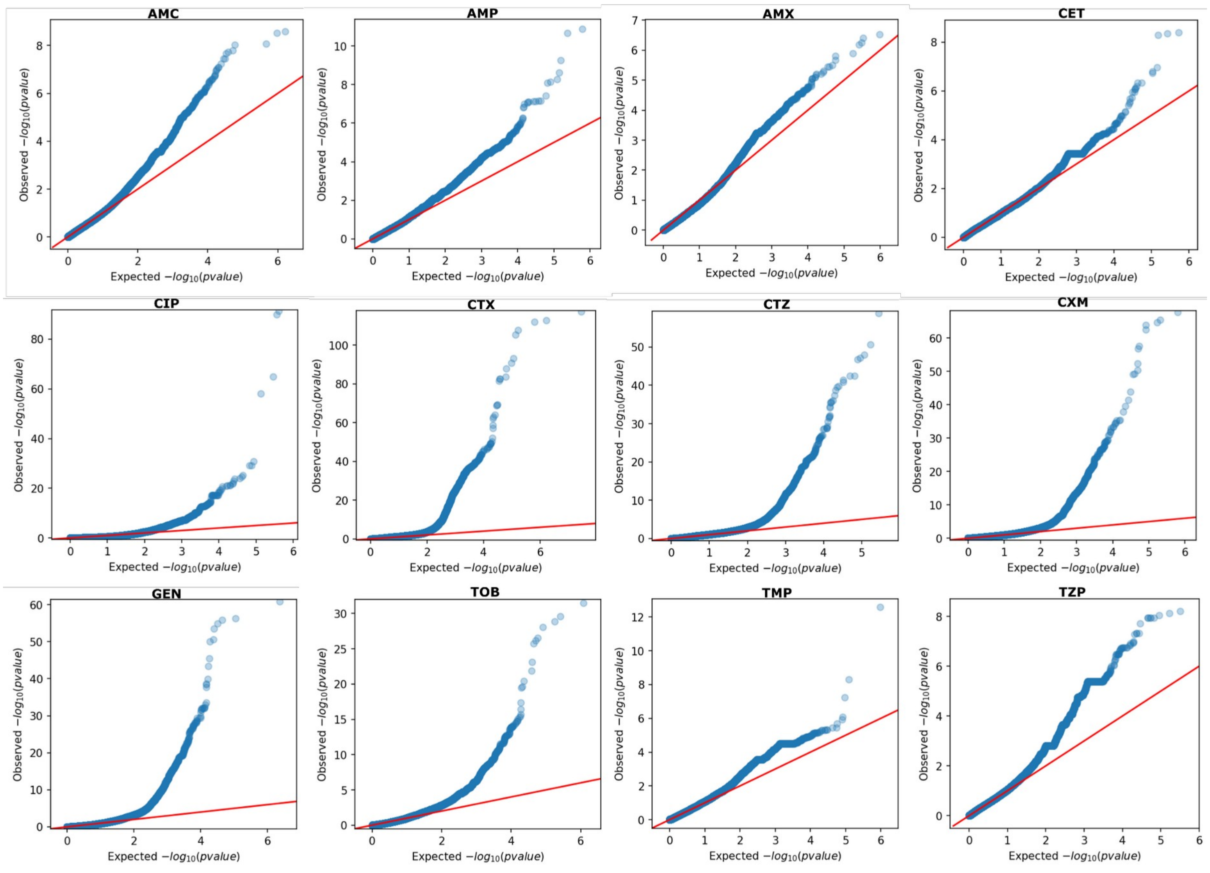


**Supplementary Figure 5**. **QQ plots comparing expected versus observed –$\log_{10}$(p-value) distributions.** Whole genome-wide association analyses were conducted between whole genome variants and resistance phenotypes in *E. coli* for twelve antibiotics. Each subplot corresponds to a different antibiotic, with the red line representing the null hypothesis of no association (i.e., expected = observed). Deviation of points above the red line indicates potential true associations.

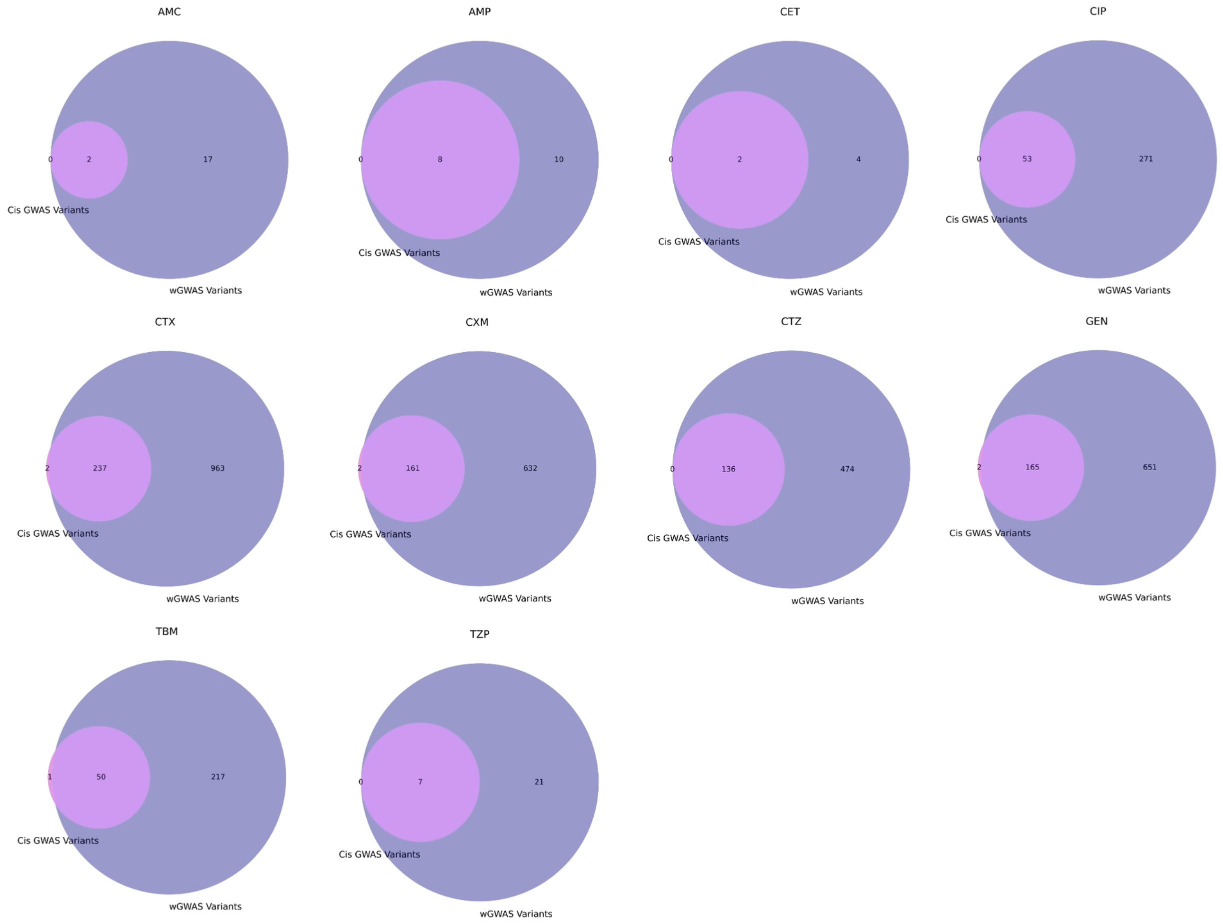


**Supplementary Figure 6: Resistance-associated variants identified by cis GWAS and whole-genome GWAS (wGWAS) in *E. coli***. Each plot illustrates the number of unique and overlapping significant variants associated with antibiotic resistance, as identified by Cis GWAS (pink) and wGWAS (purple) for the tested antibiotics. The numbers indicate the count of significant variants detected exclusively by one method or shared between both approaches.

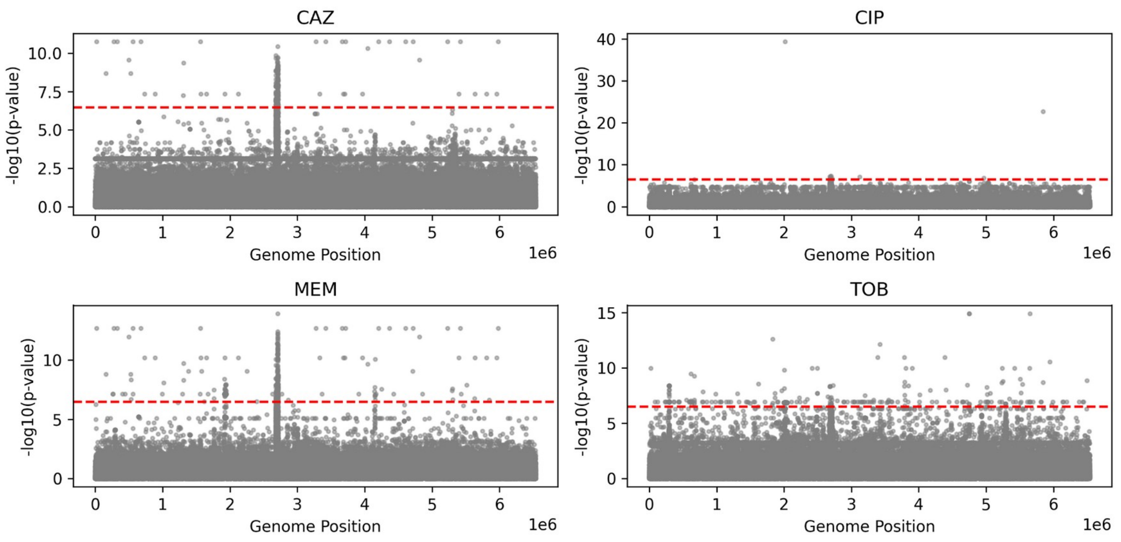


**Supplementary Figure 7: Manhattan plots for whole-genome GWAS in *P. aeruginosa.*** Each plot shows the association between an antibiotic and the resistance phenotype. Each grey dot represents a SNP, with the x-axis indicating the variant's position in the genome. The y-axis shows the p-values corrected for multiple testing. Variants above the red dashed line are considered statistically significant.

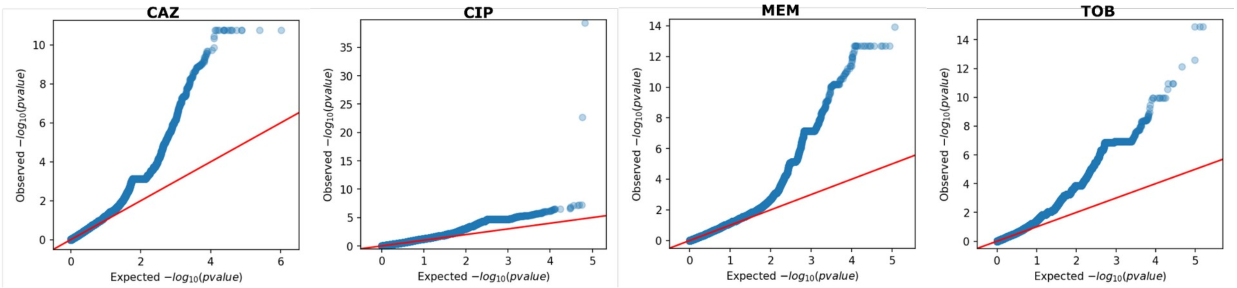


**Supplementary Figure 8**: **QQ plots displaying the relationship between expected and observed –log$_{10}$(p-value) distributions.** Whole genome-wide association analyses were conducted between whole genome variants and resistance phenotypes in *P. aeruginosa* for four antibiotics. Each plot corresponds to a different antibiotic. The red diagonal line represents the null hypothesis of no association. Upward deviations from the red line indicate inflation of observed p-values, suggesting potential genetic associations with resistance to the respective antibiotic.

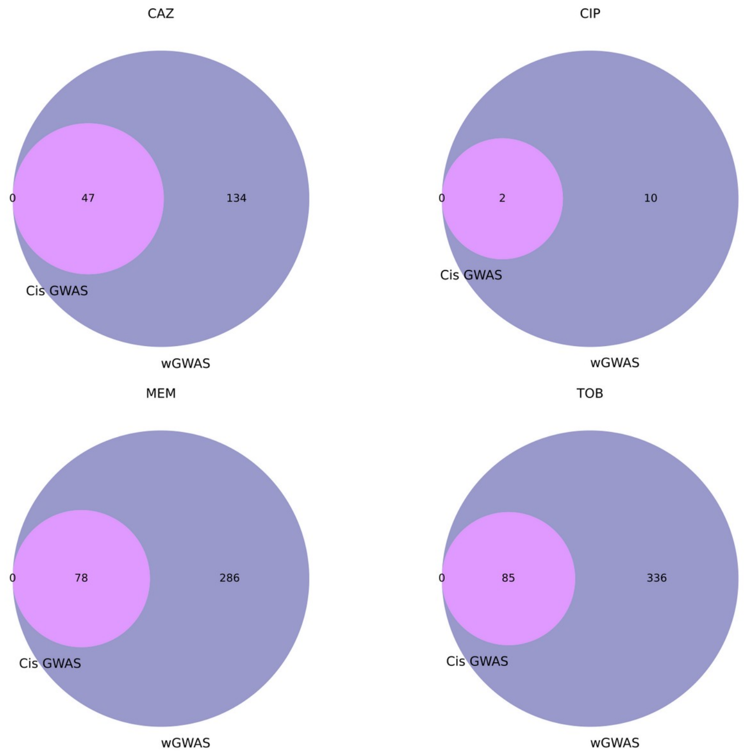


**Supplementary Figure 9: Resistance-associated variants identified by cis GWAS and whole-genome GWAS (wGWAS) in *P. aeruginosa***. Each plot illustrates the number of unique and overlapping significant variants associated with antibiotic resistance, as identified by Cis GWAS (pink) and wGWAS (purple) for the tested antibiotics. The numbers indicate the count of significant variants detected exclusively by one method or shared between both approaches.
